# Short-term effectiveness of HIV care coordination among persons with recent HIV diagnosis or history of poor HIV outcomes

**DOI:** 10.1101/311597

**Authors:** Denis Nash, McKaylee M. Robertson, Kate Penrose, Stephanie Chamberlin, Rebekkah S. Robbins, Sarah L. Braunstein, Julie E. Myers, Bisrat Abraham, Sarah Kulkarni, Levi Waldron, Bruce Levin, Mary K. Irvine

## Abstract

The New York City HIV Care Coordination Program (CCP) combines multiple evidence-based strategies to support persons living with HIV (PLWH) at risk for, or with a recent history of, poor HIV outcomes. We assessed the comparative effectiveness of the CCP by merging programmatic data on CCP clients with population-based surveillance data on all New York City PLWH. A non-CCP comparison group of similar PLWH who met CCP eligibility criteria was identified using surveillance data. The CCP and non-CCP groups were matched on propensity for CCP enrollment within four baseline treatment status groups (newly diagnosed or previously diagnosed and either consistently unsuppressed, inconsistently suppressed or consistently suppressed). We compared CCP to non-CCP proportions with viral load suppression at 12-month follow-up. Among the 13,624 persons included, 15·3% were newly diagnosed; among the 84·7% previously diagnosed, 14·2% were consistently suppressed, 28·9% were inconsistently suppressed, and 41 ·6% were consistently unsuppressed in the year prior to baseline. At 12-month follow-up, 59·9% of CCP and 53·9% of non-CCP participants had viral load suppression (Relative Risk=1.11, 95%CI:1.08-1.14). Among those newly diagnosed and those consistently unsuppressed at baseline, the relative risk of viral load suppression in the CCP versus non-CCP participants was 1.15 (95%CI:1.09-1.23) and 1.32 (95%CI:1.23-1.42), respectively. CCP exposure shows benefits over no CCP exposure for persons newly diagnosed or consistently unsuppressed, but not for persons suppressed in the year prior to baseline. We recommend more targeted case finding for CCP enrollment and increased attention to viral load suppression maintenance.

## Introduction

HIV care continuum outcomes remain suboptimal throughout the US (1). In 2013, an estimated 1·1 million persons were living with HIV infection; an estimated 87% were diagnosed; and 55% of persons diagnosed achieved viral load suppression (VLS) (1). Efforts aimed at controlling the domestic HIV epidemic will require integrated medical and social support approaches in order to extend the benefits of HIV treatment to the large numbers of persons living with HIV (PLWH) who to date have not been able to achieve and sustain VLS (2).

A first step toward strengthening the HIV care continuum is to address immediate barriers to adherence and to improve short-term outcomes among PLWH who are under-engaged with HIV medical care and treatment. HIV affects vulnerable and marginalized populations, and PLWH have higher rates of mental illness, substance use disorders, and unstable housing, all serious and often co-occurring barriers to achieving desired HIV outcomes (3, 4). Research has shown that services specifically addressing these barriers can facilitate improved outcomes along the care continuum (5–10). However, few studies have examined whether comprehensive and coordinated HIV care management programs that address both medical and social/supportive service needs are more effective at promoting VLS compared to usual-care approaches for those with known barriers to HIV care and treatment adherence (5, 11–14).

We conducted an observational study to assess the comparative effectiveness of a comprehensive HIV care coordination intervention on VLS outcomes among persons in New York City (NYC) with documented barriers to HIV care and treatment engagement (5).

## Materials and Methods

### Intervention Description

In December 2009, with Ryan White Part A funding, the NYC Department of Health and Mental Hygiene (DOHMH) launched the HIV Care Coordination Program (CCP) to support persons at high risk for, or with a recent history of, poor HIV care outcomes. The CCP combines various evidence-based programmatic elements including case management, patient navigation, directly observed therapy (DOT), structured health promotion in home/field visits, and outreach to assist patients in accessing medical care and support services, such as mental health treatment, substance use treatment, and housing assistance. The intensity and focus of these services can be tailored to meet individual needs and circumstances. The intervention has previously been described, and CCP materials are available on the DOHMH <u>website</u> (11, 15).

Importantly, the CCP was rolled out purely as a service program, with no randomization or contemporaneous control/comparison group. Early assessments of CCP outcomes used individuals as their own historical controls (i.e., pre-post) (5, 11, 15). While these assessments offered preliminary evidence suggestive of program effectiveness, they could not distinguish program effects from secular improvements in VLS in NYC during the same time period.(16) Focusing on the last viral load (VL) in a 12-month follow-up period, this study aimed to compare VLS proportions between CCP clients and demographically and clinically similar PLWH who, during the same time period, were eligible for but did not enroll in the CCP (“non-CCP PLWH”).

### Data Sources

We retrospectively constructed an observational cohort of persons enrolled and not enrolled in the CCP by merging provider-reported programmatic data with data from the longitudinal population-based NYC HIV Surveillance Registry (“the Registry”). The Registry contains demographic and laboratory information on all diagnoses of HIV (since 2000) and AIDS (since 1981) reported in NYC, with the addition of comprehensive HIV-related laboratory reporting (including all CD4-lymphocyte [CD4] and VL test results) starting in 2005. The Registry does not contain direct measures of primary care or HIV treatment status (17). Vital status information is updated through regular matches with local and national death data. Data on CCP client enrollments were drawn from the DOHMH Electronic System for HIV/AIDS Reporting and Evaluation (eSHARE), a secure, Web-based, named programmatic reporting system.

In eSHARE, we identified all persons who enrolled in the CCP from December 1, 2009 to March 31, 2013. Using data reported to the Registry as of September 30, 2014, we identified all persons who were diagnosed with HIV as of March 31, 2013, were living 12 months after diagnosis, were at least 18 years old as of March 31, 2013, and had at least one CD4 or VL result dated between December 1, 2007 and March 31, 2013. To ensure adequate outcome observation time, we excluded CCP clients who died within 12 months of program enrollment (n=279).

This study was approved by the institutional review boards at The City University of New York and the New York City Department of Health and Mental Hygiene. For these secondary analyses of de-identified data, we received a waiver for informed consent under 45 CFR 46.116(d)(2).

### ‘non-CCP PLWH’ comparison group

We constructed a non-CCP comparison group of PLWH who were similar to CCP enrollees in four steps. First, through the NYC Registry match, we identified PLWH who were not enrolled in the CCP but met broad clinical eligibility criteria for CCP enrollment at one or more times (CCP eligibility window) during December 1, 2009 to March 31, 2013. Second, we assigned eligible non-CCP PLWH pseudo-enrollment dates falling within their windows of CCP eligibility. Third, we limited to PLWH with evidence of recent NYC HIV medical care. Finally, we matched CCP enrollees to non-CCP PLWH according to baseline treatment status, enrollment/pseudo-enrollment dates and propensity for enrollment in the CCP.

### Step 1. Identify persons meeting broad CCP eligibility criteria

Using information from the Registry, we identified persons as eligible for enrollment in the CCP if they were 1) *newly diagnosed* with HIV from December 1, 2008 to March 31, 2013; 2) *out of medical care*, defined as lacking CD4 and VL laboratory monitoring for any nine-month post-diagnosis period during December 1, 2007 to March 31, 2013; 3) *treatment naïve*, defined as ever having a CD4 count <200 reported as of March 31, 2013, but not initiating antiretroviral treatment [ART] (never having a ≥1 −log drop in VL within 3 months, or an unsuppressed VL [>200 copies/μL] followed by a suppressed VL [≤200 copies/μL]) as of the date of a CD4 count <200 (17); 4) *exhibiting poor ART adherence* as of March 31, 2012, defined as not achieving VLS or not having any VL tests reported in the first 12 months after ART initiation (a ≥1 −log drop in VL within 3 months, or an unsuppressed VL followed by a suppressed VL) (17); 5) *experiencing viral rebound* (a suppressed VL followed by 2 consecutive unsuppressed VL tests in the 12 months following the suppressed VL, from December 1, 2007 to March 31, 2013); or 6) *registering a high VL* (≥10,000 copies/μL) from December 1, 2008 to March 31, 2013.

### Step 2. Assign eligible persons in the non-CCP PLWH comparison group a pseudo-enrollment date

Non-CCP PLWH who met any of the eligibility criteria were assigned an *eligibility window* (Supplemental Table 1), or a range of time between December 2009 and March 2013 (the CCP enrollment period), during which they met the above CCP eligibility criteria. For example, persons were considered eligible as “newly diagnosed” during the 12 months following diagnosis. Persons could be assigned multiple eligibility windows based on qualifying for the CCP via multiple Registry criteria and/or qualifying under the same criterion multiple times. To identify a start of follow-up for each member of the comparison group (i.e., time zero from which to prospectively assess outcomes in comparison to those in the CCP group), we randomly assigned each non-CCP PLWH a pseudo-enrollment date that fell within one of their eligibility windows. Further, to control for secular trends in VLS, pseudo-enrollment dates were assigned with probabilities such that their temporal distribution matched that of the enrollment dates among CCP enrollees (i.e., frequency matching). For persons who died, eligibility windows ended at least 12 months prior to the date of death, to ensure 12 months for outcome observation following the pseudo-enrollment date.

### Step 3. Identify NYC medical care recipients in the non-CCP PLWH group

After pseudo-enrollment dates were assigned, we restricted the eligible pool to persons who had at least one valid CD4 or VL test reported to the Registry in the 24 months after the pseudo-enrollment date. We required one laboratory test to identify persons accessing HIV medical care in NYC after the pseudo-enrollment date, as CCP enrollment and service initiation entails connection to NYC HIV medical care (18).

### Step 4. Propensity Model and Match

After constructing this eligible non-CCP-enrolled population, we prepared to match persons in the non-CCP PLWH comparison group to those in the CCP group using baseline treatment status, propensity scores, and enrollment/pseudo-enrollment dates. Given that 12-month VLS outcomes would be expected to differ by baseline treatment status/engagement, we created four mutually exclusive baseline treatment status groups: 1) newly diagnosed (in the 12 months prior to enrollment/pseudo-enrollment date), 2) consistently suppressed (≥2 VLs ≥90 days apart, and all VLs ≤200 copies/μL, in the 12 months prior to enrollment/pseudo-enrollment date), 3) consistently *un*suppressed (all VLs reported >200 copies/μL or no VLs reported in the 12 months prior to enrollment/pseudo-enrollment), or 4) inconsistently suppressed (at least 1 VL ≤200 copies/μL, but *not all* VLs ≤200 copies/μL, in the 12 months prior to enrollment/pseudo-enrollment).

We used logistic regression to estimate the propensity for enrollment in the CCP within each of the above 4 groups. We combined two baseline treatment status groups, groups 3 and 4 for one propensity score model, because we hypothesized the propensity of enrollment in the CCP would be influenced by the same potential confounders for these two groups; when we examined the models separately, the effect estimates from the pooled model did not differ from the effect estimates from the individual models. We report the results for the pooled model because we were able to match more participants. Subsequent matching occurred within each of the four groups.

For the three propensity score models, we started with a model that included all of our *a priori* hypothesized and measured confounders and used backward selection to identify the model with the lowest value of Akaike’s Information Criterion (AIC). The variables that we suspected were confounders of the relationship between CCP enrollment and the VLS outcome were sex, race/ethnicity, age at enrollment/pseudo-enrollment, country of birth, HIV transmission risk, year of diagnosis, baseline VL, baseline CD4, successful linkage to HIV care within three months of diagnosis, presence of an AIDS diagnosis within one year of HIV diagnosis, number of VL laboratory tests reported in the year prior to enrollment/pseudo-enrollment, residential ZIP code at enrollment/pseudo-enrollment, HIV prevalence and poverty level within ZIP code at enrollment/pseudo-enrollment, and interaction terms for baseline CD4 and baseline VL, baseline CD4 and race, sex and risk, and year of diagnosis and risk. Per the American Community Survey, poverty level within the residential ZIP code at enrollment/pseudo-enrollment was classified as high (poverty greater than the median poverty level for a given year of enrollment/pseudo-enrollment) versus low. HIV prevalence was based on aggregated NYC HIV surveillance data for the ZIP code by year of enrollment/pseudo-enrollment, and was classified as high (prevalence greater than the median HIV prevalence for a given year) versus low.

Within each of the four baseline treatment status groups, we matched on propensity scores and enrollment/pseudo-enrollment dates (± 3 months). We used a 1:1 ‘greedy’ match technique, and the match algorithm proceeded sequentially from 8 to 1 decimal places of the propensity score (19, 20). We considered a standardized difference of ≥0·1 to indicate an imbalance in the measured confounders between the CCP and non-CCP groups (21). The final match included no imbalances ≥0·1.

### Outcome Definition and Care Coordination Program Effectiveness Estimate

The VLS outcome was based on the last VL laboratory result reported to the Registry in the 12 months following the enrollment/pseudo-enrollment date, and was dichotomized as ≤200 or >200 copies/μL. Persons with no VL in the Registry for the entire 12-month follow-up period were classified as not having VLS. While rare, 1 ·3% (87/6,812) of the CCP and 5·4% (353/6,812) of the non-CCP PLWH were missing a VL result. We used a GEE model with binary error distribution and identity link to estimate the difference in proportion of CCP and non-CCP participants with VLS, accounting for the matched-pairs design. The arithmetic difference was expressed in percentage point units. We used a GEE model with binary error distribution and log link to estimate the relative effect of the CCP on VLS, accounting for the matching. Absolute differences and relative risks were estimated with the GENMOD procedure in SAS version 9·3 (SAS Institute, Cary, N.C.) The absolute and relative effect of the CCP was estimated for each of the four baseline treatment status groups.

## Results

From December 1, 2009 to March 31, 2013, a total of 7,337 persons enrolled in the CCP; of those, 7,058 (96·2%) were still living 12 months after enrollment. Of the 62,828 non-enrolled PLWH who were eligible for enrollment in the CCP, 91·9% (57,746) were assigned a pseudo-enrollment date; 74·8% (46,997) had an HIV-related laboratory test in NYC in the two years following their pseudo-enrollment date; and 10·8% (6,812) were matched to a CCP enrollee, allowing us to include 96·5% (6,812/7,058) of CCP clients eligible for this analysis (Figure 1).

**Figure 1.**
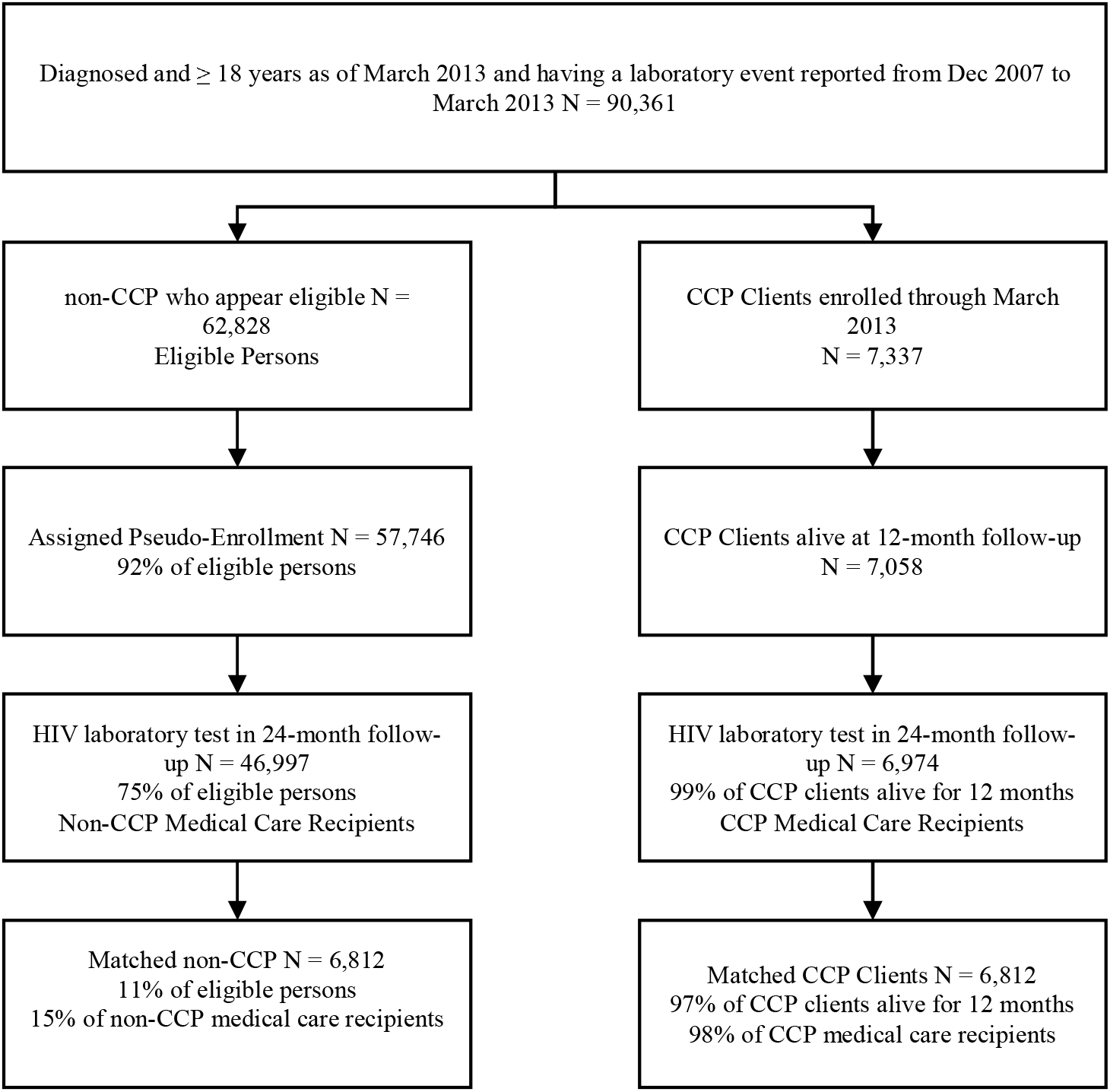
Flow Chart for Study Inclusion in the Care Coordination and non-Care Coordination Comparison Group Analysis, New York City, 2009-2013

Prior to propensity score matching, the CCP and non-CCP groups differed substantially by measured demographic and clinical characteristics (Table 1). After propensity score matching, the CCP and non-CCP groups were similar on all measured characteristics (Table 1). In terms of baseline treatment status, 15·3% were newly diagnosed in the year prior to enrollment/pseudo-enrollment, and among the 84·7% who were previously diagnosed, 14·2% were consistently virally suppressed, 28·9% were inconsistently suppressed, and 41·6% were consistently *un*suppressed in the year prior to enrollment.

**Table 1.** Pre and Post-Match Demographic and Clinical Characteristics of Care Coordination and non-Care Coordination Program Persons - New York City, 2009-2013.

|  | Pre-Match |  |  | Post-Match |  |  |
| --- | --- | --- | --- | --- | --- | --- |
|  | Total | non-CCP | CCP | Total | non-CCP | CCP |
| Total | 53,971 (100.0) | 46,997 (100.0) | 6,974 (100.0) | 13,624 (100.0) | 6,812 (100.0) | 6,812 (100.0) |
| Sex |  |  |  |  |  |  |
| Male | 38,380 (71.1) | 33,920 (72.2) | 4,460 (64.0) | 8,750 (64.2) | 4,379 (64.3) | 4,371 (64.2) |
| Female | 15,591 (28.9) | 13,077 (27.8) | 2,514 (36.0) | 4,874 (35.8) | 2,433 (35.7) | 2,441 (35.8) |
| Race/Ethnicity |  |  |  |  |  |  |
| Black | 25,646 (47.5) | 21,931 (46.7) | 3,715 (53.3) | 7,248 (53.2) | 3,603 (52.9) | 3,645 (53.5) |
| Hispanic | 17,339 (32.1) | 14,682 (31.2) | 2,657 (38.1) | 5,260 (38.6) | 2,678 (39.3) | 2,582 (37.9) |
| White | 9,766 (18.1) | 9,338 (19.9) | 428 (6.1) | 820 (6.0) | 395 (5.8) | 425 (6.2) |
| Other/Unknown | 1,220 (2.2) | 1,046 (2.3) | 174 (2.5) | 296 (2.2) | 136 (2.0) | 160 (2.3) |
| Age Category (years) |  |  |  |  |  |  |
| ≤24 | 3,551 (6.6) | 2,999 (6.4) | 552 (7.9) | 1,070 (7.9) | 534 (7.8) | 536 (7.9) |
| 25-44 | 21,931 (40.6) | 18,985 (40.4) | 2,946 (42.2) | 5,820 (42.7) | 2,944 (43.2) | 2,876 (42.2) |
| 45-64 | 26,540 (49.2) | 23,288 (49.6) | 3,252 (46.6) | 6,301 (46.2) | 3,113 (45.7) | 3,188 (46.8) |
| 65+ | 1,949 (3.6) | 1,725 (3.7) | 224 (3.2) | 433 (3.2) | 221 (3.2) | 212 (3.1) |
| Transmission Risk |  |  |  |  |  |  |
| Men who have sex with men | 20,440 (37.9) | 18,410 (39.2) | 2,030 (29.1) | 3,986 (29.3) | 1,995 (29.3) | 1,991 (29.2) |
| Injection drug use history | 8,916 (16.5) | 7,433 (15.8) | 1,483 (21.3) | 2,916 (21.4) | 1,474 (21.6) | 1,442 (21.2) |
| Heterosexual | 11,832 (21.9) | 9,930 (21.1) | 1,902 (27.3) | 3,678 (27.0) | 1,836 (27.0) | 1,842 (27.0) |
| Other/unknown | 12,783 (23.7) | 11,224 (23.9) | 1,559 (22.4) | 3,044 (22.3) | 1,507 (22.1) | 1,537 (22.6) |
| Country of Birth |  |  |  |  |  |  |
| US/US dependency | 35,563 (65.9) | 30,932 (65.8) | 4,631 (66.4) | 9,127 (67.0) | 4,571 (67.1) | 4,556 (66.9) |
| Foreign born | 9,764 (18.1) | 8,160 (17.4) | 1,604 (23.0) | 3,022 (22.2) | 1,498 (22.0) | 1,524 (22.4) |
| Unknown | 8,644 (16.0) | 7,905 (16.8) | 739 (10.6) | 1,475 (10.8) | 743 (10.9) | 732 (10.7) |
| Year of HIV Diagnosis |  |  |  |  |  |  |
| Prior 1995 | 9,989 (18.5) | 8,689 (18.5) | 1,300 (18.6) | 2,529 (18.6) | 1,258 (18.5) | 1,271 (18.7) |
| 1995-1999 | 9,976 (18.5) | 8,738 (18.6) | 1,238 (17.8) | 2,396 (17.6) | 1,181 (17.3) | 1,215 (17.8) |
| 2000-2004 | 14,684 (27.2) | 12,862 (27.4) | 1,822 (26.1) | 3,621 (26.6) | 1,813 (26.6) | 1,808 (26.5) |
| 2005-2009 | 11,549 (21.4) | 10,120 (21.5) | 1,429 (20.5) | 2,858 (21.0) | 1,464 (21.5) | 1,394 (20.5) |
| 2010-2013 | 7,773 (14.4) | 6,588 (14.0) | 1,185 (17.0) | 2,220 (16.3) | 1,096 (16.1) | 1,124 (16.5) |
| Baseline Viral Load (copies/μL) |  |  |  |  |  |  |
| >1500 | 17,634 (32.7) | 13,820 (29.4) | 3,814 (54.7) | 7,374 (54.1) | 3,666 (53.8) | 3,708 (54.4) |
| >200-1499 | 6,749 (12.5) | 6,039 (12.8) | 710 (10.2) | 1,423 (10.4) | 713 (10.5) | 710 (10.4) |
| ≤200 | 21,108 (39.1) | 18,927 (40.3) | 2,181 (31.3) | 4,266 (31.3) | 2,141 (31.4) | 2,125 (31.2) |
| No viral load | 8,480 (15.7) | 8,211 (17.5) | 269 (3.9) | 561 (4.1) | 292 (4.3) | 269 (3.9) |
| Baseline CD4 Count (cells/μL) |  |  |  |  |  |  |
| <200 | 8,758 (16.2) | 6,466 (13.8) | 2,292 (32.9) | 4,335 (31.8) | 2,135 (31.3) | 2,200 (32.3) |
| 200-349 | 9,182 (17.0) | 7,684 (16.3) | 1,498 (21.5) | 2,953 (21.7) | 1,486 (21.8) | 1,467 (21.5) |
| 350 - 499 | 9,668 (17.9) | 8,469 (18.0) | 1,199 (17.2) | 2,416 (17.7) | 1,228 (18.0) | 1,188 (17.4) |
| 500+ | 18,392 (34.1) | 16,678 (35.5) | 1,714 (24.6) | 3,356 (24.6) | 1,668 (24.5) | 1,688 (24.8) |
| No CD4 | 7,971 (14.8) | 7,700 (16.4) | 271 (3.9) | 564 (4.1) | 295 (4.3) | 269 (3.9) |
| Initiated Care ≤91 Days |  |  |  |  |  |  |
| No | 37,412 (69.3) | 32,701 (69.6) | 4,711 (67.6) | 9,243 (67.8) | 4,615 (67.7) | 4,628 (67.9) |
| Yes | 16,559 (30.7) | 14,296 (30.4) | 2,263 (32.4) | 4,381 (32.2) | 2,197 (32.3) | 2,184 (32.1) |
| Baseline HIV Prevalence & Poverty |  |  |  |  |  |  |
| High poverty & high prevalence | 28,689 (53.2) | 23,985 (51.0) | 4,704 (67.5) | 9,240 (67.8) | 4,647 (68.2) | 4,593 (67.4) |
| Low poverty & high prevalence | 12,473 (23.1) | 11,227 (23.9) | 1,246 (17.9) | 2,412 (17.7) | 1,188 (17.4) | 1,224 (18.0) |
| High poverty & low prevalence | 1,740 (3.2) | 1,495 (3.2) | 245 (3.5) | 459 (3.4) | 227 (3.3) | 232 (3.4) |
| Low poverty & low prevalence | 7,003 (13.0) | 6,327 (13.5) | 676 (9.7) | 1,311 (9.6) | 650 (9.5) | 661 (9.7) |
| Unknown | 4,066 (7.5) | 3,963 (8.4) | 103 (1.5) | 202 (1.5) | 100 (1.5) | 102 (1.5) |
| Number of Viral Load Labs in year prior to enrollment |  |  |  |  |  |  |
| 0 VL labs | 8,480 (15.7) | 8,211 (17.5) | 269 (3.9) | 561 (4.1) | 292 (4.3) | 269 (3.9) |
| 1-3 VL labs | 28,923 (53.6) | 25,169 (53.6) | 3,754 (53.8) | 7,404 (54.3) | 3,729 (54.7) | 3,675 (53.9) |
| 4+ VL labs | 16,568 (30.7) | 13,617 (29.0) | 2,951 (42.3) | 5,659 (41.5) | 2,791 (41.0) | 2,868 (42.1) |
| Baseline Treatment Status |  |  |  |  |  |  |
| Newly diagnosed<sup>1</sup> | 7,682 (14.2) | 6,590 (14.0) | 1,092 (15.7) | 2,088 (15.3) | 1,044 (15.3) | 1,044 (15.3) |
| Consistently suppressed<sup>2</sup> | 5,939 (11.0) | 4,917 (10.5) | 1,022 (14.7) | 1,934 (14.2) | 967 (14.2) | 967 (14.2) |
| Consistently *unsuppressed*<sup>3</sup> | 21,631 (40.1) | 18,742 (39.9) | 2,889 (41.4) | 5,666 (41.6) | 2,833 (41.6) | 2,833 (41.6) |
| Inconsistently suppressed<sup>4</sup> | 18,719 (34.7) | 16,748 (35.6) | 1,971 (28.3) | 3,936 (28.9) | 1,968 (28.9) | 1,968 (28.9) |

1. Newly diagnosed within 12 months of enrollment or pseudo-enrollment
2. Consistently suppressed: at least 2 VLs at least 90 days apart and all VLs < 200 copies/mL in 12 months prior to enrollment or pseudo-enrollment
3. Consistently *unsuppressed*: All labs reported >200 in 12 months prior to enrollment or pseudo-enrollment. This includes persons missing VL labs. Persons with 1 lab >200 would be in this group.
4. Inconsistently suppressed: Previously diagnosed and not in groups (2) and (3) above. Persons with 1 lab ≤200 would be in this group.

Only 31·3% were virally suppressed at the time of last reported lab measurement prior to enrollment/pseudo-enrollment, and the median CD4 count was 314 cells/μL (IQR 150-506), with 31·8% having CD4<200 cells/μL.

By twelve months after enrollment/pseudo-enrollment, 59·9% of the CCP group and 53·9% of the non-CCP comparison group were suppressed on their most recent VL test (Table 2). The proportion of persons with VLS differed by baseline treatment status. Two baseline treatment status groups showed significantly higher CCP versus non-CCP VLS in the follow-up year: newly diagnosed PLWH (73·3% versus 63·3%, respectively; Absolute Difference as a percentage (AD): 9·96 (95%CI 6·09-13·83); Relative Risk (RR):1·15 (95%CI 1·09-1·23); and PLWH who were previously diagnosed and consistently *un*suppressed in the year prior to enrollment/pseudo-enrollment (42·5% vs. 32·1%, respectively; AD:10·41 (95%CI 7·94-12·89); RR:1·32 (95%CI 1·23-1·42)). No differences in VLS were observed among persons with consistent VLS or with inconsistent VLS in the year prior to enrollment/pseudo-enrollment (91·7% vs. 90·6%; AD_consistent VLS_=1·14 (95%CI −1·42-3·69); RR_consistent VLS_=1·01 (95%CI 0·98-1·04); and 62·2% vs. 62·3%; AD_inconsistent VLS_=−0·10 (95%CI: −3·08-2·88); RR_inconsistent VLS_=0·99 (95%CI: 0·95-1·05), respectively).

**Table 2.** Relative and Absolute Difference for having Viral Load Suppression at 12-Month Post Measure (Care Coordination Program Versus non-Care Coordination Program Persons) – New York City, 2009-2013.

|  | Overall<br>(N = 14,060) |  |  | CCP | non-CCP | Arithmetic<br>Difference in<br>Percentage Points<br>(95% Confidence<br>Intervals (CI)) | Relative Risk<br>(95% CI) | P-Value<sup>2</sup> |
| --- | --- | --- | --- | --- | --- | --- | --- | --- |
|  | Total | %<br>VLS<br>Post<sup>1</sup> | N<br>Denominator<br>(for CCP or<br>non-CCP) | %<br>VLS<br>Post<sup>1</sup> | %<br>VLS<br>Post<sup>1</sup> |  |  |  |
| **Overall** | **13,624** | **56.9** | **6,812** | **59.9** | **53.9** | 5.99 (4.49, 7.49) | **1.11 (1.08, 1.14)** | <0.001 |
| **Baseline Treatment Status** |  |  |  |  |  |  |  |  |
| Newly diagnosed<sup>3</sup> | 2,088 | 68.3 | 1,044 | 73.3 | 63.3 | 9.96 (6.09, 13.83) | **1.15 (1.09, 1.23)** | <0.001 |
| Consistently suppressed<sup>4</sup> | 1,934 | 91.2 | 967 | 91.7 | 90.6 | 1.14 (-1.42, 3.69) | 1.01 (0.98, 1.04) | 0.38 |
| Consistently unsuppressed<sup>5</sup> | 5,666 | 37.3 | 2,833 | 42.5 | 32.1 | 10.41 (7.94, 12.89) | **1.32 (1.23, 1.42)** | <0.001 |
| Inconsistently suppressed<sup>6</sup> | 3,936 | 62.3 | 1,968 | 62.2 | 62.3 | -0.10 (-3.08, 2.88) | 0.99 (0.95, 1.05) | 0.95 |

CCP: Care Coordination Program, CI: Confidence Intervals, VLS: Viral Load Suppression
1. Proportion with the last viral load ( $\leq 200$ ) in the 12 months after enrollment or pseudo-enrollment. Persons missing a VL are considered to have unsuppressed VL ( $>200$ )
2. P-values for the arithmetic difference in proportions and their relative risks did not differ to the number of decimal places shown
3. Newly diagnosed within 12 months of enrollment or pseudo-enrollment
4. Consistently suppressed: at least 2 VLs at least 90 days apart and all VLs $< 200$ copies/mL in 12 months prior to enrollment or pseudo-enrollment
5. Consistently *unsuppressed*: All labs reported $>200$ in 12 months prior to enrollment or pseudo-enrollment. This includes persons missing VL labs. Persons with 1 lab $>200$ would be in this group.
6. Inconsistently suppressed: Previously diagnosed and not in groups (2) and (3) above. Persons with 1 lab $\leq 200$ would be in this group.

## Discussion

Using a surveillance-based method for comparison group selection in an observational effectiveness study, we found New York’s Ryan White CCP intervention to have a significant positive effect on VLS for newly diagnosed persons and for previously diagnosed persons who were consistently *un*suppressed in the year prior to enrollment. Previously, we have shown a consistent CCP effect using a single-group pre-post assessment (i.e., individuals serving as their own controls), with the proportion of CCP-enrollees with VLS increasing from 32.3% in the year prior to enrollment to 50.9% in the year after enrollment.(5, 11) However, the single-group pre-post design could not isolate program effects from secular improvements in VLS (i.e., annual citywide improvements in VLS that occurred in tandem with population-based HIV treatment strategies).(5, 11, 22, 23) A strength of our contemporaneous comparison group approach is that it accounts for these secular trends in VLS; these data suggest the CCP may help PLWH with initial hurdles to ART access and adherence, and point to the potential value of targeting the intervention to these individuals who have not previously achieved VLS. However, twelve months after CCP enrollment, there remained substantial room for improvement in VLS among those CCP participants who were *un*suppressed throughout the year prior to enrollment, as only 42.5% had achieved VLS on their last VL measurement.

There was a high prevalence of advanced HIV disease in the cohort (31·8% had CD4<200 cells/uL), with a substantial proportion of persons not suppressed on their last VL measure in the 12-month follow-up period (4·31%). This likely reflects the persistence of major barriers to care and treatment engagement in our study population. For the sample of 7,058 CCP clients alive at 12-month follow-up, we previously described the baseline prevalence of psychosocial barriers, including unstable housing (22·1%), low mental health functioning (30·0%), and self-reported recent hard drug use (15·1%); half of CCP clients (50·4%, or 3,556) had documentation of at least one of these barriers at the time of enrollment, and only 24·4% of the 3,556 (or 38·6% of the 2,250 with a follow-up assessment) were documented as having none of those barriers at last assessment in the 12 months post-enrollment (5). Thus, housing, mental health and/or drug-related barriers persisted for ≥61·4% (and up to 75·6%) of clients presenting with those barriers at CCP enrollment.

Both CCP and non-CCP groups with inconsistent viral suppression at baseline were already engaged in HIV care and treatment to some degree, and the lack of a significant CCP effect in these groups may suggest that the CCP is less suited to helping people stay adherent. This should be examined in an assessment of longer-term (durable) VLS. Additionally, the NYC epidemic has shown substantial improvements in VLS over time (24), likely driven by advances in ART, treatment guideline expansion, and a robust local and state system of medical and non-medical services for PLWH. High-quality ‘usual care’ would tend to mute intervention effects in all baseline treatment status groups.

Strengths of our study include the use of a population-based data source to rigorously derive a contemporaneous observational comparison group, which was large enough to identify a non-CCP PLWH match (similar with regard to measured factors) for 96·5% of the CCP sample. Additionally, deriving outcome information for both CCP and non-CCP recipients from the HIV Registry ensured that it was highly complete across the cohort and over time, regardless of care location or changes in care provider within NYC. Finally, our method’s explicit attention to matching on enrollment/pseudo-enrollment dates controlled for secular trends of increasing VLS in NYC over time.

Limitations of our study include those related to observational studies, such as the possibility of uncontrolled or poorly controlled confounding. Since our propensity models were limited to variables available in the NYC HIV Surveillance Registry. Additionally, we were not able to describe or account for services or service delivery models received in the non-CCP comparison group. Finally, our results may not be fully generalizable to Ryan White clients in settings outside NYC.

## Conclusions

New York’s Ryan White CCP intervention has shown a positive short-term effect on VLS among newly diagnosed PLWH and those who were consistently virally *un*suppressed in the year prior to the start of follow-up, suggesting the program may be effective at helping with initial hurdles to ART access and adherence. However, the absence of an effect among persons with any suppression in the year prior to enrollment indicates the program may be less effective at helping people stay adherent. CCP efforts in NYC could better target for enrollment those newly diagnosed and consistently *un*suppressed, and increase emphasis on maintenance of viral suppression. Future studies should include assessment of longer-term outcomes, including sustained viral suppression, in this population and others with known barriers to HIV care and treatment.

## Acknowledgements

The authors would like to acknowledge Care Coordination Program providers and clients, for their input at different stages of this work and their invaluable contributions to real-world program implementation and evaluation.

## Disclaimer

The findings and conclusions in this report are those of the authors and do not necessarily represent the official position of the National Institutes of Health.

